# metaboprep v2: Broadening the application of the *metaboprep* beyond metabolomics

**DOI:** 10.1101/2025.11.11.687831

**Authors:** Nicholas Sunderland, David A Hughes, Mathew A Lee, Alec McKinlay, Nicholas J Timpson, Laura J Corbin

## Abstract

High-throughput multiplex assays for metabolomics and proteomics offer opportunities for biomarker discovery and disease stratification in epidemiological research. The complexity of these datasets requires robust, standardized and transparent preprocessing workflows to ensure reproducibility and comparability across studies. We present an updated and enhanced version of the *metaboprep* R package. Originally designed for metabolomics data, it has now been extended to support proteomics datasets from platforms such as Olink® and SomaScan®. This release introduces a user-friendly, modular, object-oriented architecture using R’s S7 system, enabling improved input format flexibility, streamlined report generation and increased compatibility with other third-party tools. The updated pipeline is structured in three parts: data import, filtering and summary, and output generation. This structure provides a reproducible yet customizable framework for pre-analysis data preparation with utility across multiple omics platforms and particular value in supporting multi-cohort epidemiological research.

**Availability and implementation:** The *metaboprep* package is implemented in R and freely available at: https://github.com/MRCIEU/metaboprep.

## Introduction

In epidemiology, the high-throughput measurement of large numbers of variables capturing substantial portions of variation in specific domains of biological variation (“omics”) has opened a range of analytical applications including prediction of disease risk, subtyping and classification of disease, biomarker discovery and mechanism dissection (Babu and Snyder, 2023). In recognition of the demand for increased consistency and transparency in the preanalytical processing of metabolomics data across cohorts and studies, we previously published the R package *metaboprep* (Hughes *et al*., 2022). This package was designed to enable the extraction of metabolomics data from different platforms and for the processing, summary and preparation of those data for subsequent statistical analysis within a standardized and reproducible workflow.

As technologies enabling the simultaneous measurement of multiple circulating proteins become increasingly used in epidemiological studies, a similar demand for standardized workflows has arisen in this field (Bowser and Robinson, 2023; Babu and Snyder, 2023). Proteomics data generated by high-throughput assays share many characteristics in common with similarly produced metabolomics data – a high number of variables or features, a range of data distributions, variable detection limits and the possible influence of technical covariates (e.g. batch effects). In this context, we have seen *metaboprep* being useful in pre-processing proteomics data derived by both Olink, using their Olink Target® platform (Goulding *et al*.; Bull *et al*., 2024), and by SomaLogic, using their SomaScan® assay (Goudswaard *et al*., 2023).

Here we describe an update to our established metabolomics pre-analysis platform (*metaboprep*), including an extension to its omics data scope. While retaining the original purpose of the package, this update to *metaboprep* (v2.0) allows a single transparent pipeline for preprocessing metabolomics data with flexibility for the user to define their own criteria for sample and feature filtering, but with updates designed to fulfil two principal objectives: (1) improved flexibility regarding input data types and formats; and (2) improved user experience and reporting.

### Toolbox

#### Pipeline design

*metaboprep* is an R package designed to standardize the steps involved in preparing large metabolomics and proteomics datasets for statistical analysis. The updated package (v2.0) retains the functionality of the previous pipeline with functions now accessible within discrete customizable modules, along with flexible data import and export workflows. This includes compatibility with community standards (COMETS Analytics (Temprosa *et al*., 2022)) and use with existing analytical software (MetaboAnalyst (Pang *et al*., 2024)). Modules can be run in isolation or combined into an end-to-end pipeline resulting in filtered datasets and data summary reporting. Data transformations, sample and feature exclusions, and pre-processing metrics, are stored within a central *Metaboprep* R S7 class object (Vaughan *et al*., 2024). This object can be used in various workflows in a way that feels familiar to R users, particularly those accustomed to *dplyr* (Wickham, Hadley *et al*., 2023) or *tidyr* (Wickham, Hadley *et al*., 2024) syntax. An overview of the overall pre-processing pipeline and underlying modules is presented in **Figure 1**.

**Figure 1.**
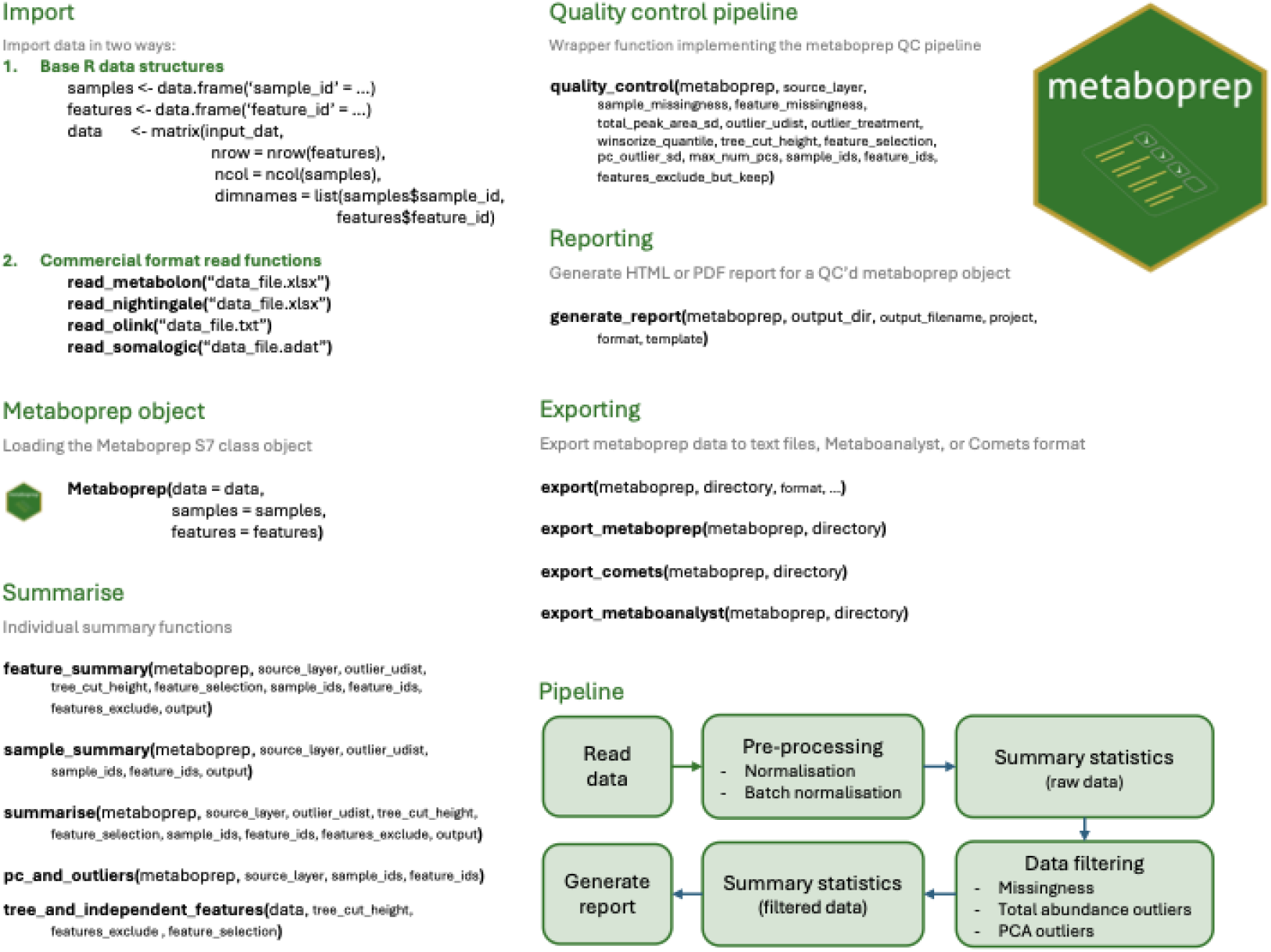
An overview ‘cheat sheet’ of the *metaboprep v2*.*0* R package functions and pre-processing pipeline.

#### Implementation

*metaboprep v2*.*0* is implemented using R (version 4.5.1) (R Core Team, 2025) and is dependent upon R version 4.1.0 or greater. Code is available from https://githubhttps://github.com/MRCIEU/metaboprep along with an associated GitHub Pages website (https://mrcieu.github.io/metaboprep/), providing a walkthrough of the package. Example datasets are available within the package for users to test and explore the utility of *metaboprep v2*.*0*. To enhance structure, extensibility, and user experience, *metaboprep v2*.*0* leverages the modern S7 object-oriented system in R. Central to this design is the ‘Metaboprep’ class - an S7 object serving as a structured container for omics data, including arrays and data frames of feature measurements, sample and feature metadata, summary statistics, and exclusion flags. The S7 framework enables robust class property validation with rule-based checks, such as ensuring alignment between sample and feature IDs and associated metadata, thereby maintaining data integrity throughout the pipeline (Vaughan *et al*., 2024).

A key feature of the ‘Metaboprep’ class is its layered data architecture, which stores multiple versions of the dataset (e.g., raw input, filtered data, and transformed data) as separate layers within a three-dimensional numerical array. This approach preserves the full processing history while allowing easy access and comparison across stages, promoting reproducibility and auditability of analyses. Moreover, all processing functions in *metaboprep v2*.*0* accept and return ‘Metaboprep’ objects, enforcing standardized workflows and consistent handling of data, and propagation of metadata. This design facilitates modular, extensible, and transparent data processing aligned with tidy software principles and reproducible research standards. To ensure backward compatibility with metaboprep v1.0, we provide the function *run_metaboprep1()*, which reproduces the original routine and accepts the legacy parameter file format.

#### Modularity

*Metaboprep v2*.*0* is designed with modularity and flexibility at its core to accommodate diverse data sources and user needs. The pipeline is organized into three distinct parts: (1) data import; (2) summary and filtering; and (3) output generation (including report creation). This segmentation improves usability and makes debugging easier. The modular structure allows users to run individual components independently or as part of an integrated workflow.

The pipeline begins with a data preparation phase that supports reading from commercial provider formats such as Nightingale Health (CoreMetabolomics), Metabolon, Inc. (Global Discovery Panel), Olink® (Olink Explore) and SomaLogic (SomaScan). The format of the standard deliverables from these companies is known to change overtime and whilst the import functions can handle the common current and legacy formats, the ability for users to input their own custom datasets through use of simple base R data structures (data frames and matrices) provides some degree of future proofing. The *read_[provider]()* functions handle data extraction from often complex Microsoft Excel and custom data formats into a list of base R data types: a numeric matrix of abundance data and data frames of sample and feature metadata. The only core requirement of the sample and feature data frames is that they contain a column named “*sample_id*” and “*feature_id*”, respectively.

The modularization of the import architecture, an improvement on the metaboprep v1.0, supports rapid adaptation to emerging platforms and evolving file specifications, futureproofing *metaboprep v2*.*0* against evolving data sources. Example (simulated) datasets emulating the structure of common provider outputs and associated ‘read in’ functions are included (https://github.com/MRCIEU/metaboprep/tree/master/inst/extdata). This allows users to trial the package without proprietary data. Example (simulated) datasets currently included as part of the package and based on data releases received by us from providers include: Metabolon, Inc. (Global Discovery Panel) v1 (pre-October 2020) and v2 (October 2020 onwards); Nightingale Health (CoreMetabolomics) v1 (pre-April 2020) and v2 (April 2020 onwards). In addition, example datasets have been generated based on code and datasets provided in GitHub repositories of Olink (https://github.com/Olink-Proteomics/OlinkRPackage) and SomaLogic (https://github.com/SomaLogic).

The ‘Metaboprep’ class is then created, with its constructor taking the data matrix and sample and feature metadata data frames. By default, the input data is entered into the object’s layered 3D data matrix (array) at index “input” along the third dimension. Validation checks are conducted at instantiation to ensure matching feature and sample identifiers in the data and enforcing correct order. Pre-analysis processing is often necessary, for example batch normalization procedures. Calling functions such as *batch_normalise()* on the ‘Metaboprep’ object inserts another layer along the object’s data array third dimension which can then be specified as the source data for subsequent operations such as the pre-processing pipeline.

At the center of *metaboprep* functionality is the pre-processing routine, called with the *quality_control()* function. This is a wrapper around underlying summarization, outlier analysis and filtering modules, that if needed can be called individually to explore the data and assist in debugging. Several arguments provide the user with flexibility in selecting samples or features to include, through the “*sample_ids*” and “*feature_ids*” arguments. There is also the possibility to exclude certain features from the outlier and filtering steps whilst retaining them in the resulting data, through use of the “*features_exclude_but_keep”* argument – this can be useful in datasets that contain derived ratio measures (Nightingale Health) or xenobiotics with high degrees of missingness (Metabolon, Inc.). The *quality_control()* function adds a further data layer along the ‘Metaboprep’ object’s data array third dimension called “qc”, representing the cleaned data ready for analysis, as well as populating the object’s “exclusion” field with a list of excluded samples and features and the reason for their exclusion.

#### Report generation

*Metaboprep v2*.*0* includes optional report generation capabilities that provide detailed summaries of the pre-processing and filtering steps that have been applied to the data. These reports facilitate transparency and reproducibility by documenting key metrics such as missingness rates, outlier detection, sample exclusions and data transformations at each processing stage. The output formats are flexible, with options to generate comprehensive HTML reports for review as well as PDF files suitable for sharing with collaborators and including in publications. An example report is included in **Supplementary Data S1**.

#### Documentation and examples

Comprehensive documentation for *metaboprep v2*.*0* is provided through multiple channels to support users of all levels. A detailed README on the GitHub repository offers step-by-step instructions for running the pipeline. To further enhance user experience, *metaboprep v2*.*0* leverages *pkgdown* to generate a user-friendly, searchable website that organizes the package documentation, reference manuals and vignettes in one accessible location (https://mrcieu.github.io/metaboprep/). This online resource offers guided aids, including tutorials and examples, helping users to quickly understand and apply the package functions. The vignettes included with the package provide practical walkthroughs and use cases.

## Conclusion

*Metaboprep v2*.*0* provides a flexible, standardized, and reproducible framework for preprocessing multi-omics data, including metabolomics and proteomics. Leveraging modern S7 object-oriented design, it ensures data integrity and transparent tracking of processing steps through layered data storage. The package also generates fully documented, platform-agnostic summary reports for supplementary materials. With improved input flexibility, modular pipeline components, and compatibility with diverse platforms, this update to *metaboprep* enhances usability and supports consistent workflows for epidemiological studies.

## Code availability

The *metaboprep* R package, along with installation instructions and source code, is available on GitHub (https://github.com/MRCIEU/metaboprep). Precompiled binaries for common platforms are also available via the MRC-IEU R Universe, enabling easy installation using install.packages(“metaboprep”, repos = c(“https://mrcieu.r-universe.dev“, “https://cloud.r-project.org“)).

## Funding

NS is funded by the GW4-CAT Wellcome PhD programme. LC, NJT and NS work in a Medical Research Council (UKRI) funded unit (MC_UU_00032/1 and MC_UU_00032/3). AM is supported by a CRUK PhD studentship [grant number C18281/A30905]. MAL is supported by funding from IIG_FULL_2021_008, which was obtained from Wereld Kanker Onderzoek Fonds, as part of the World Cancer Research Fund International grant programme, and from INCA_15849, which was obtained from Institut National du Cancer.

Where authors are identified as personnel of the International Agency for Research on Cancer/World Health Organization, the authors alone are responsible for the views expressed in this article and they do not necessarily represent the decisions, policy, or views of the International Agency for Research on Cancer/World Health Organization.

## Figure legends

**Supplementary Data S1**. An example PDF report generated using *metaboprep v2*.*0*.

## Notes

https://github.com/MRCIEU/metaboprep

https://mrcieu.github.io/metaboprep/

### Competing Interest Statement

The authors have declared no competing interest.

