## Supplement 1 for "metaboprep v2: Broadening the application of the *metaboprep* beyond metabolomics"

### MetaboPrep Data Preparation Summary Report

09 September, 2025

**8 Power analysis** **31**

### 1 Project Information

This `metaboprep` report summarizes the data preparation steps for:

- **Project:** Project

#### 1.1 Overview

The `metaboprep` R package performs three key operations:

1. **Assessment & Summary Statistics**: Provides an initial assessment of raw metabolomics data.
2. **Data Filtering**: Applies filtering techniques to clean the dataset.
3. **Post-Filtering Assessment**: Evaluates the filtered dataset, particularly in relation to batch variables when available.

This report contains descriptive information on both raw and filtered metabolomics data for **Project**.

Please raise any issues on the GitHub issues page.

#### 1.2 Data preparation workflow:

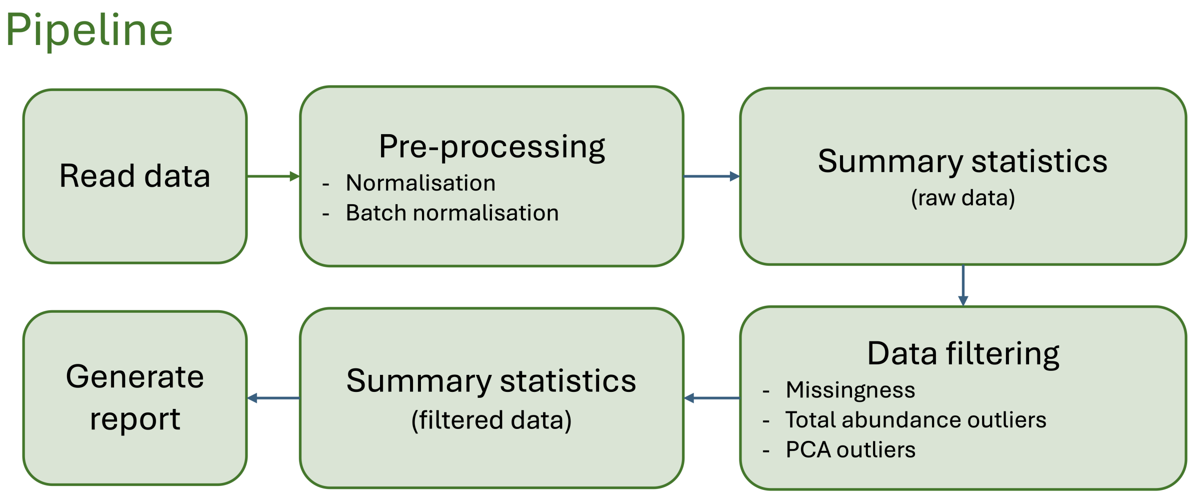

### 2 Summary of raw data

#### 2.1 Sample size of Project data set:

Table 1: Sample Size Summary

| Dataset | Samples | Features |
|---|---|---|
| INPUT | 1440 | 955 |
| QC | 1440 | 955 |

### 3 Missingness

Missingness is evaluated across samples and features using the original (input) data set.

#### 3.1 Visual structure of missingness in the raw data

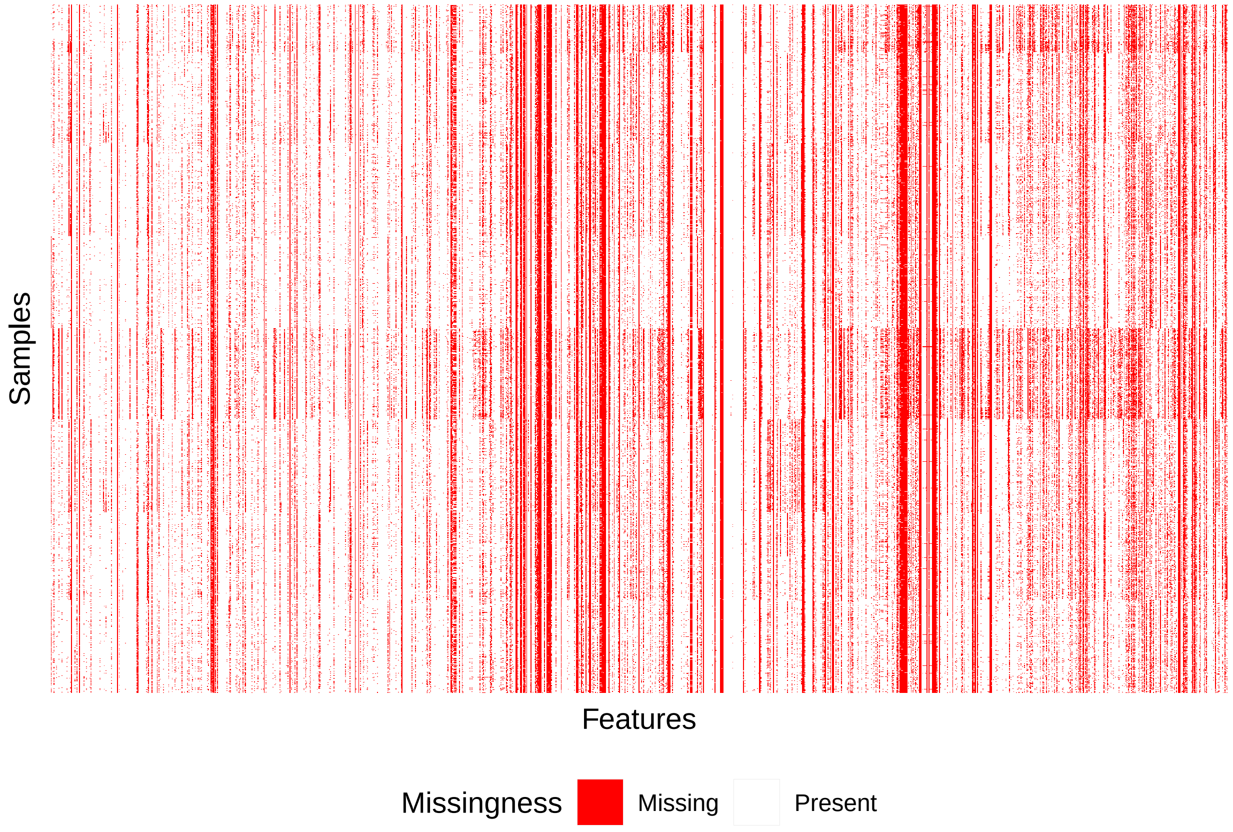

#### 3.2 Summary of sample and feature missingness

The extent of missing data in both samples and features of the dataset was evaluated - for each sample, the proportion of features with missing values, and for each feature, the proportion of samples with missing values. This can identify whether missingness is more strongly driven by poorly measured features or by problematic samples.

The distribution of missingness is summarized in:

1. histograms

2. tabulated using percentiles, and

3. estimates of study samples sizes under various levels of feature / sample missingness filtering thresholds - this helps in understanding how filtering sample and/or features with high levels of missing data would affect the dataset

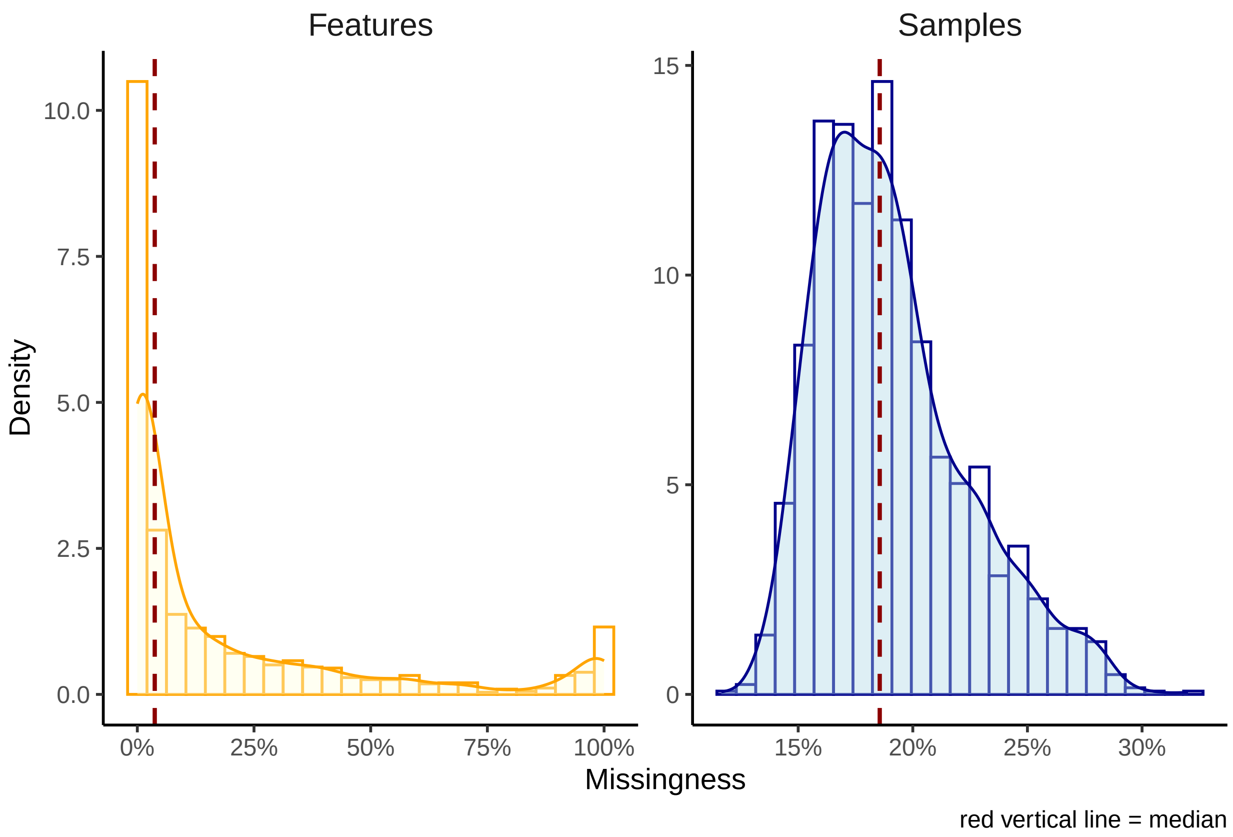

Table 2: Sample and feature missingness percentiles.

| | Percentile | Features | Samples |
|---|---|---|---|
| 0% | 0.00 | 0.000 | 0.116 |
| 25% | 0.25 | 0.001 | 0.167 |
| 50% | 0.50 | 0.037 | 0.186 |
| 75% | 0.75 | 0.263 | 0.208 |
| 100% | 1.00 | 1.000 | 0.320 |

Table 3: Estimates of samples sizes using different feature missingness thresholds.

| Feature missingness threshold | Sample size |
|---|---|
| 0% | 0 |
| 20% | 1500 |
| 40% | 1500 |
| 60% | 1500 |
| 80% | 1500 |
| 100% | 1500 |

Table 4: Estimates of samples sizes using different sample missingness thresholds.

| Sample missingness threshold | Sample size |
|---|---|
| 0% | 0 |
| 20% | 1022 |
| 40% | 1500 |
| 60% | 1500 |
| 80% | 1500 |
| 100% | 1500 |

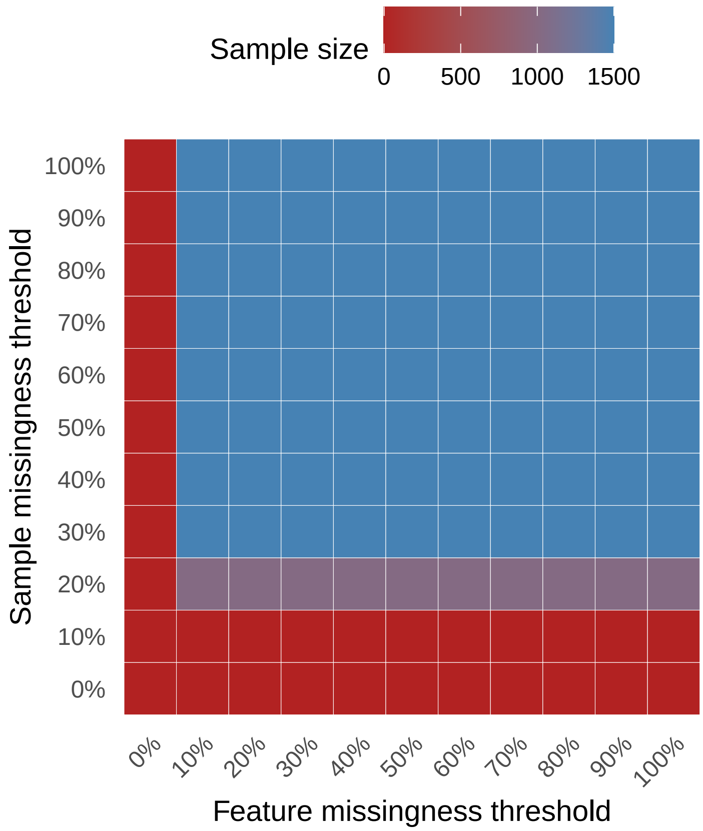
Sample size
0
500
1000
1500
100%
90%
80%
70%
60%
50%
40%
30%
20%
10%
0%
Sample missingness threshold
0%
10%
20%
30%
40%
50%
60%
70%
80%
90%
100%
Feature missingness threshold

### 4 Data Filtering

#### 4.1 Exclusion summary

Table 5: Sample and Feature Exclusions

| Reason | Count |
|---|---|
| *Features* | |
| - user excluded | 0 |
| - extreme feature missingness | 113 |
| - user defined feature missingness | 263 |
| *Samples* | |
| - user excluded | 0 |
| - extreme sample missingness | 0 |
| - user defined sample missingness | 53 |
| - user defined sample totalpeakarea | 2 |
| - user defined sample pca outlier | 5 |

*Note:*
Eight primary data filtering exclusion steps were made during the preparation of the data. User-defined thresholds indicated with an asterisk. In addition, the number of outlier datapoints modified during QC are presented.
[1] Features manually excluded
[2] Features with missingness >=80
[3] Features exclusions based on the user defined threshold of *>=20
[4] Samples manually excluded
[5] Samples with missingness >=80
[6] Samples exclusions based on user defined threshold of *>=20
[7] Samples with a total-peak-area or total-sum-abundance that is *>=5 SD from the mean.
[8] Samples that are *>=5 SD from the mean on principal components 1-to-2

#### 4.2 Metabolite or feature reduction and principal components

A data reduction was carried out to identify a list of representative features for generating a sample principal component analysis. This step reduces the level of inter-correlation in the data to ensure that the principal components are not driven by groups of correlated features.

The data reduction table presents the number of metabolites at each phase of the data reduction (Spearman's correlation distance tree cutting) analysis.

Table 6: Feature summary

| Data reduction | Count |
|---|---|
| Total metabolite count | 1331 |
| Metabolites included in data reduction | 945 |
| Number of metabolite clusters | 483 |
| Number of representative metabolites | 483 |

The following plot rerspresents principal components 1 and 2 using 945 representative metabolites. The red vertical and horizontal lines indicate the standard deviation cutoffs for identifying individual outliers. Outliers are those >=5 SD from the mean of PCs 1-2.

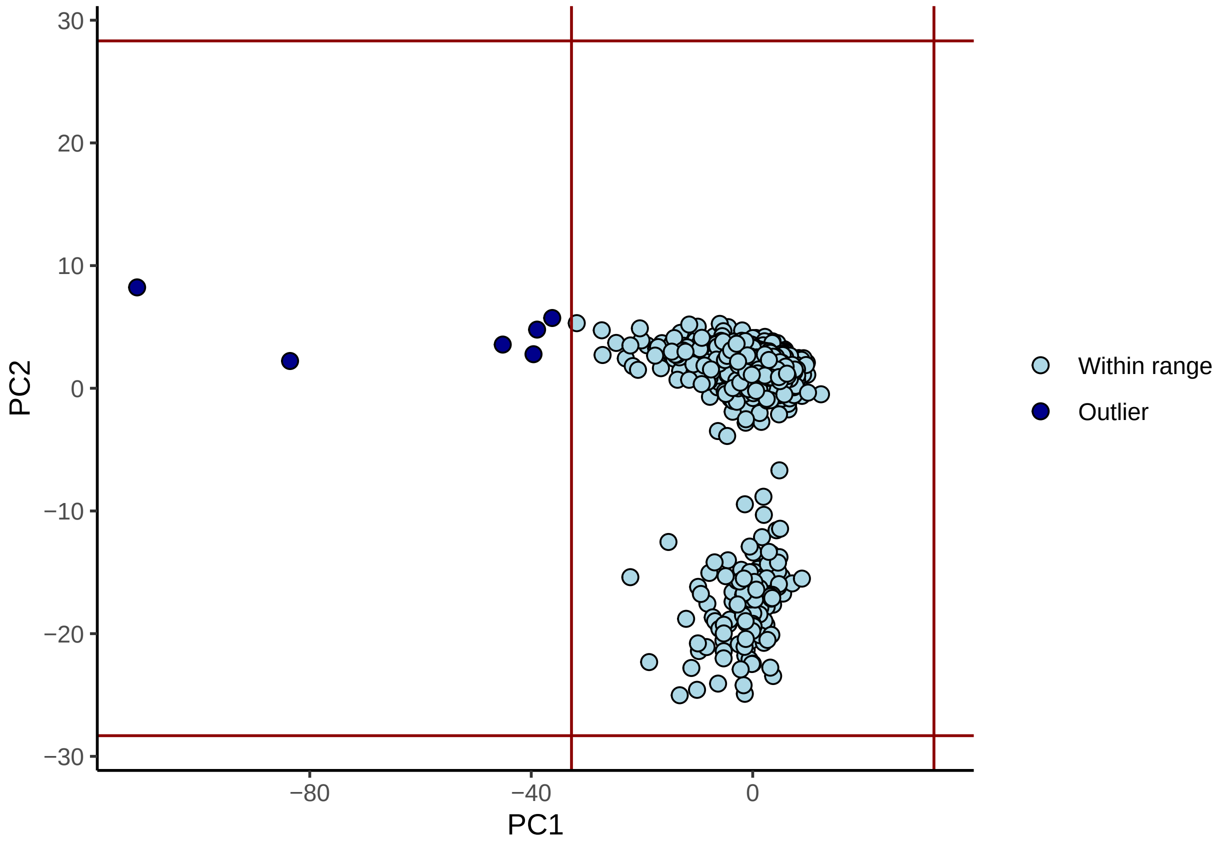

### 5 Summary of filtered data

#### 5.1 Sample size (N)

- The number of samples in data = 1440
- The number of features in data = 955

#### 5.2 Relative to the raw data

- 60 samples were filtered out, given the user's criteria.
- 376 features were filtered out, given the user's criteria.
- Please review details above and your log file for the number of features and samples excluded and why.

##### 5.2.1 Distributions for sample and feature missingness

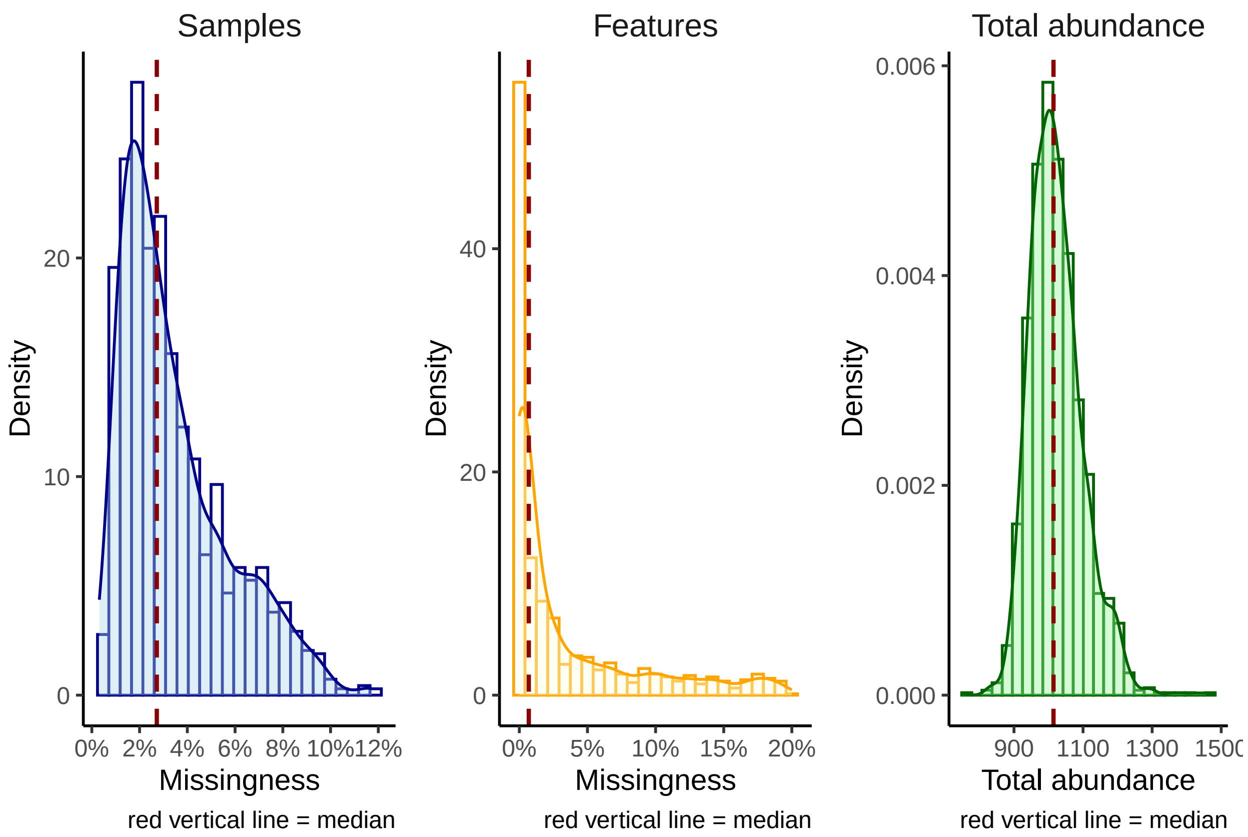

###### 5.2.2 Clustering dendrogram of representative features

Spearman's correlation distance clustering dendrogram highlighting the metabolites used as representative features in blue, the clustering tree cut height is denoted by the horizontal line.

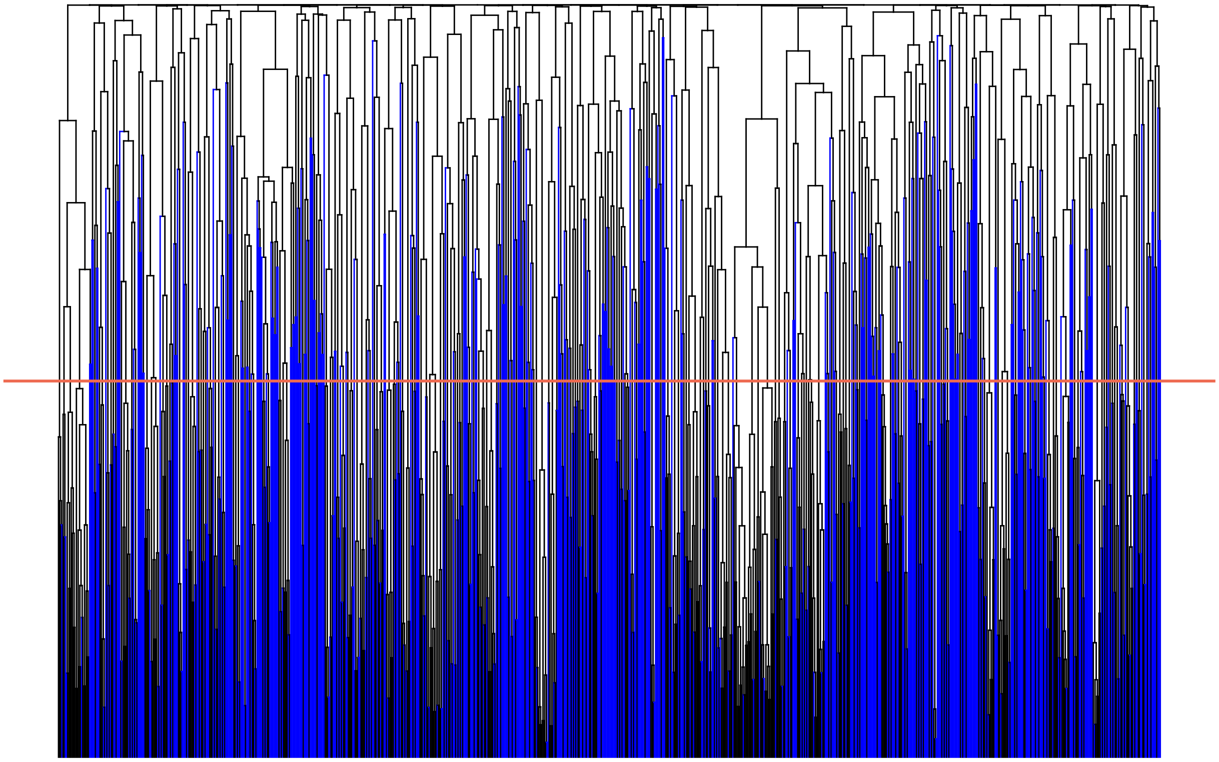

###### 5.2.3 Summary of the QC (filtered) metabolite data

The data reduction table presents the number of metabolites at each phase of the data reduction (Spearman's correlation distance tree cutting) analysis.

Table 7: QC Feature Summary

| | Count |
|---|---|
| Total metabolite count | 1331 |
| Metabolites included in data reduction | 955 |
| Number of metabolite clusters | 491 |
| Number of representative metabolites | 491 |

###### 5.2.4 Scree plot

Scree plot of the variance explained by each PC (limited to 100 for plotting) and a plot of principal component 1 and 2, as derived from the representative metabolites. The Scree plot also identifies the number of PCs estimated to be informative (vertical lines) by the Cattel's Scree Test acceleration factor (red, n = 2) and Parallel Analysis (green, n = 54).

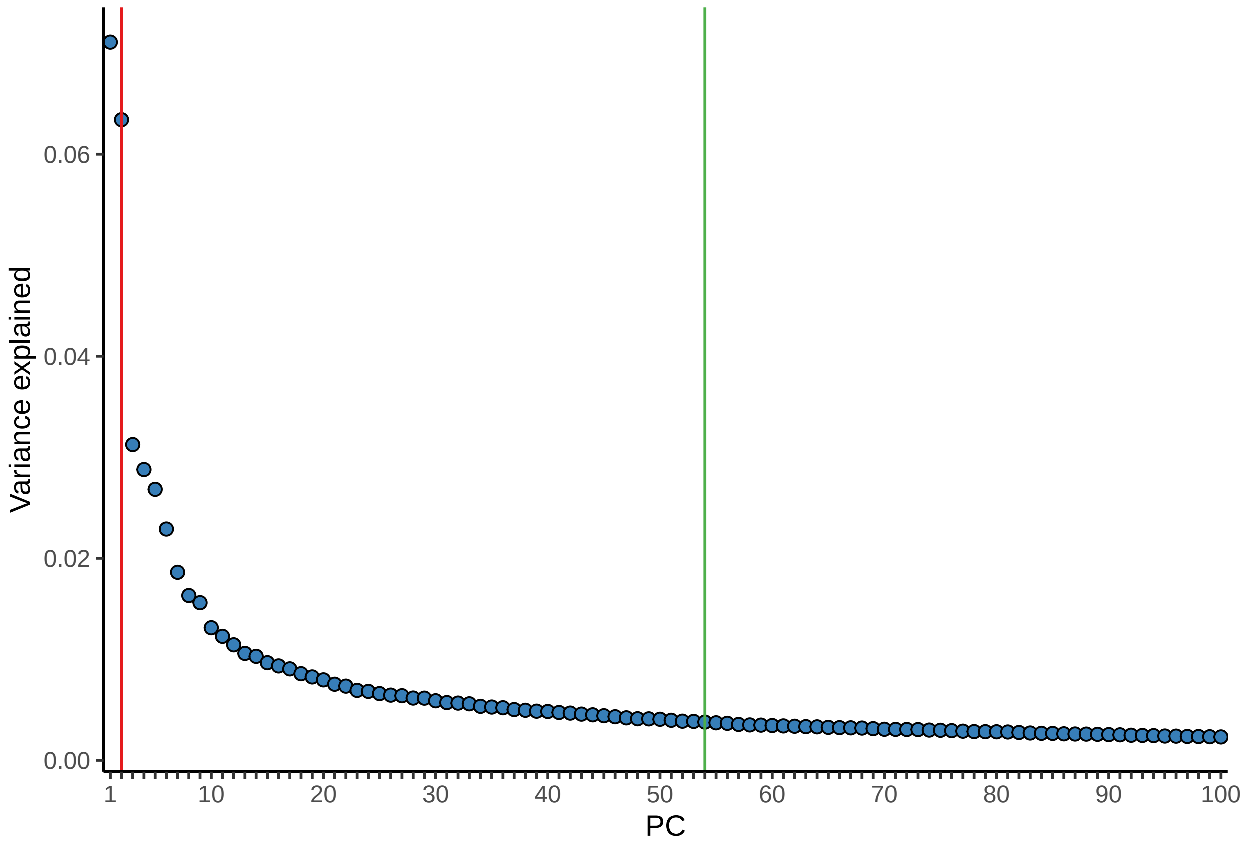

###### 5.2.5 PC plot

Individuals in the PC plot were clustered into 4 kmeans (k) clusters, using data from PC1 and PC2. The kmeans clustering and color coding is strictly there to help provide some visualization of the major axes of variation in the sample population(s).

The plot presents principal components 1 & 2 using 491 representative metabolites.

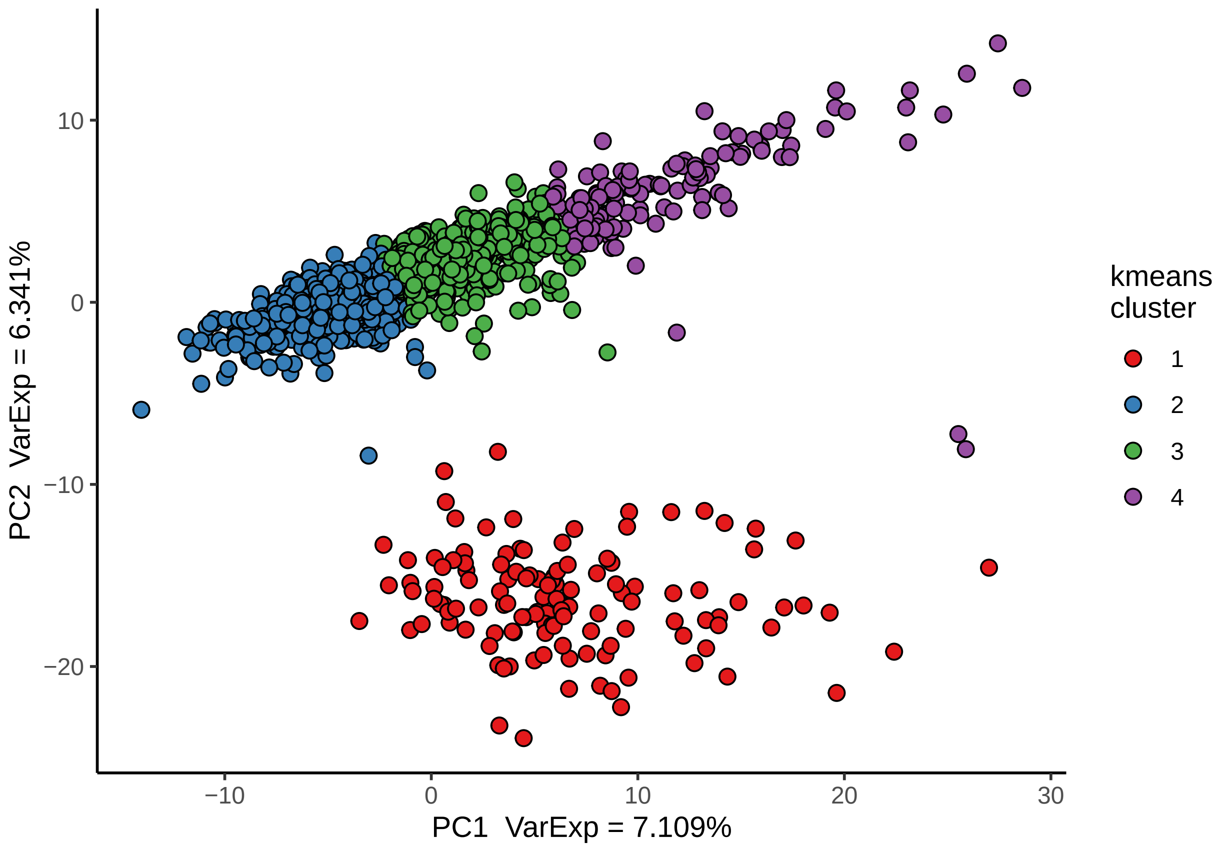

PC2 VarExp = 6.341%
PC1 VarExp = 7.109%
kmeans cluster
1
2
3
4
10
0
−10
−20
−10
0
10
20
30

#### 5.3 Structure among samples

A matrix (pairs) plot of the top five principal components including demarcations of the 3rd (grey), 4th (orange), and 5th (red) standard deviations from the mean. Samples are color coded as in the summary PC plot above using a kmeans analysis of PC1 and PC2 with a k (number of clusters) set at 4. The choice of k = 4 was not robustly chosen it was a choice of simplicity to help aid visualize variation and sample mobility across the PCs.

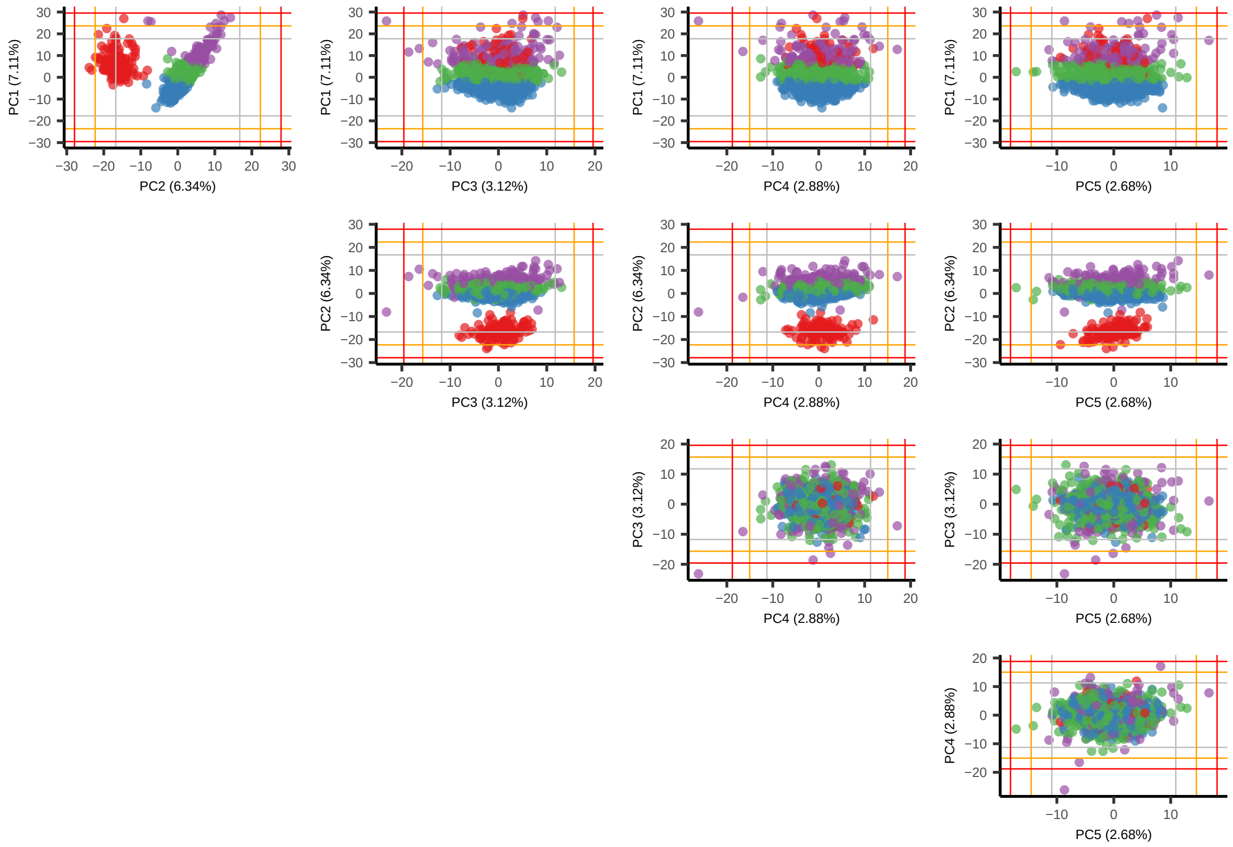

#### 5.4 Feature Distributions

##### 5.4.1 Estimates of normality: W-statistics for raw and log transformed data

Of the 1331 features in the data 376 features were excluded from this analysis because of no variation or too few observations (n < 40). Of the remaining 955 metabolite features, a total of 116 may be considered normally distributed given a Shapiro W-statistic >= 0.95.

##### 5.4.2 Distribution of W Statistics on Raw and Log10 Metabolite Abundances

Histogram plots of Shapiro W-statistics for raw and log transformed data distributions. A W-statistic value of 1 indicates the sample distribution is perfectly normal and value of 0 indicates it is perfectly uniform. Please note that log transformation of the data *may not* improve the normality of your data.

116 of the metabolites exhibit distributions that may declared normal, given a W-stat >= 0.95. In 93 instances (9.74%) of the tested metabolites the log10 data W-stat is < raw data W-stat.

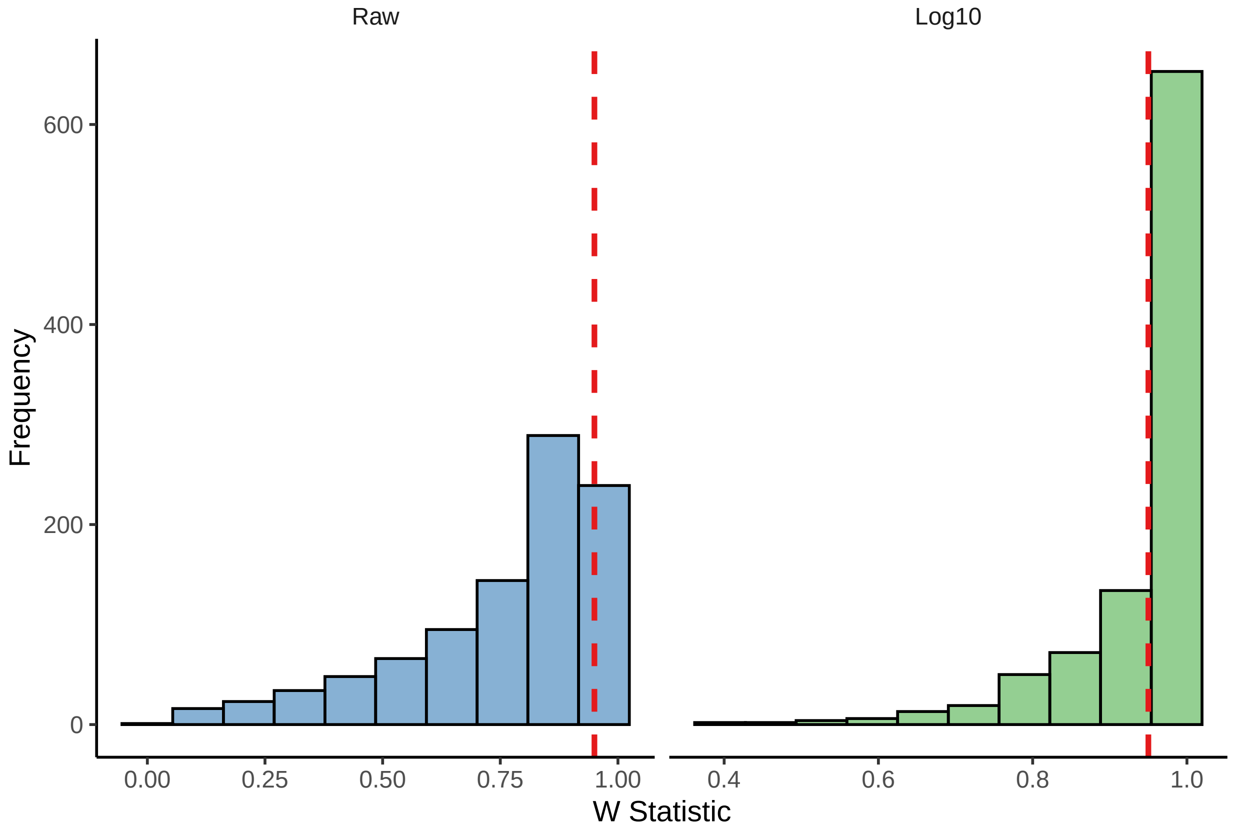

#### 5.5 Outliers

Evaluation of the number of samples and features that are outliers across the QC data. The below table presents the average number of outlier values for samples and features in the QC data set.

Table 8: Outlier Summary

| | Min. | 25th | Median | Mean | 75th | Max. |
|---|---|---|---|---|---|---|
| Features | 0 | 0 | 2 | 10.420 | 9 | 127 |
| Samples | 0 | 2 | 5 | 9.242 | 10 | 76 |

##### 5.5.1 Notes on outlying samples at each feature

There may be extreme outlying observations at individual features that have not been accounted for. You may want to:

1. Turn these observations into NAs.
2. Winsorize the data to some maximum value.
3. Rank normalize the data which will place those outliers into the top of the ranked standard normal distribution.
4. Turn these observations into NAs and then impute them along with other missing data in your data set.

### 6 Variation in filtered data by available variables

#### 6.1 Feature missingness

Feature missingness may be influenced by the features' biology or pathway classification, or the measurement methodology. The figure(s) below provides an illustrative evaluation of the proportion of *feature missigness* as a product of the variable(s) available in the raw data files.

```
-- After filtering a total of 4 feature level batch variables were identified. --
-- They are:
   LIB_ID
   PLATFORM
   SUPER_PATHWAY
   TYPE
```

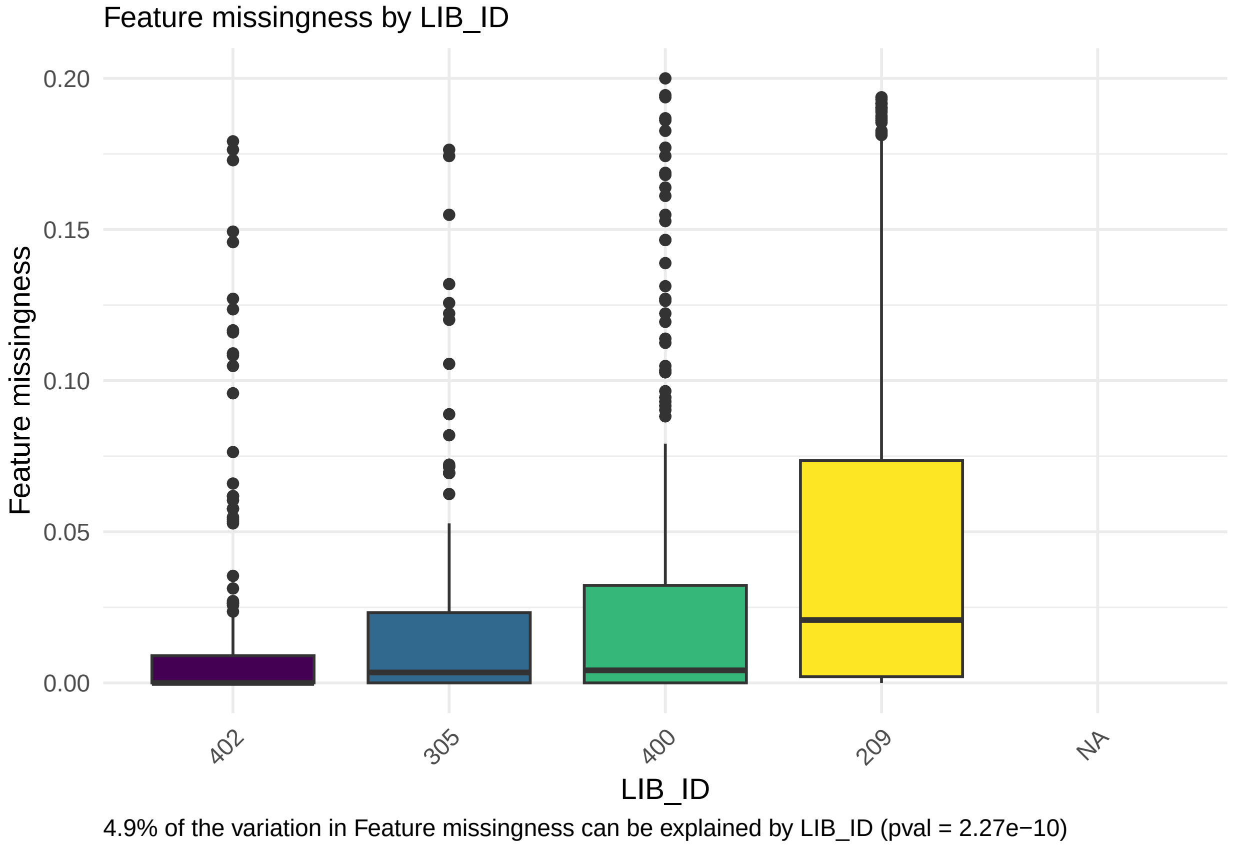

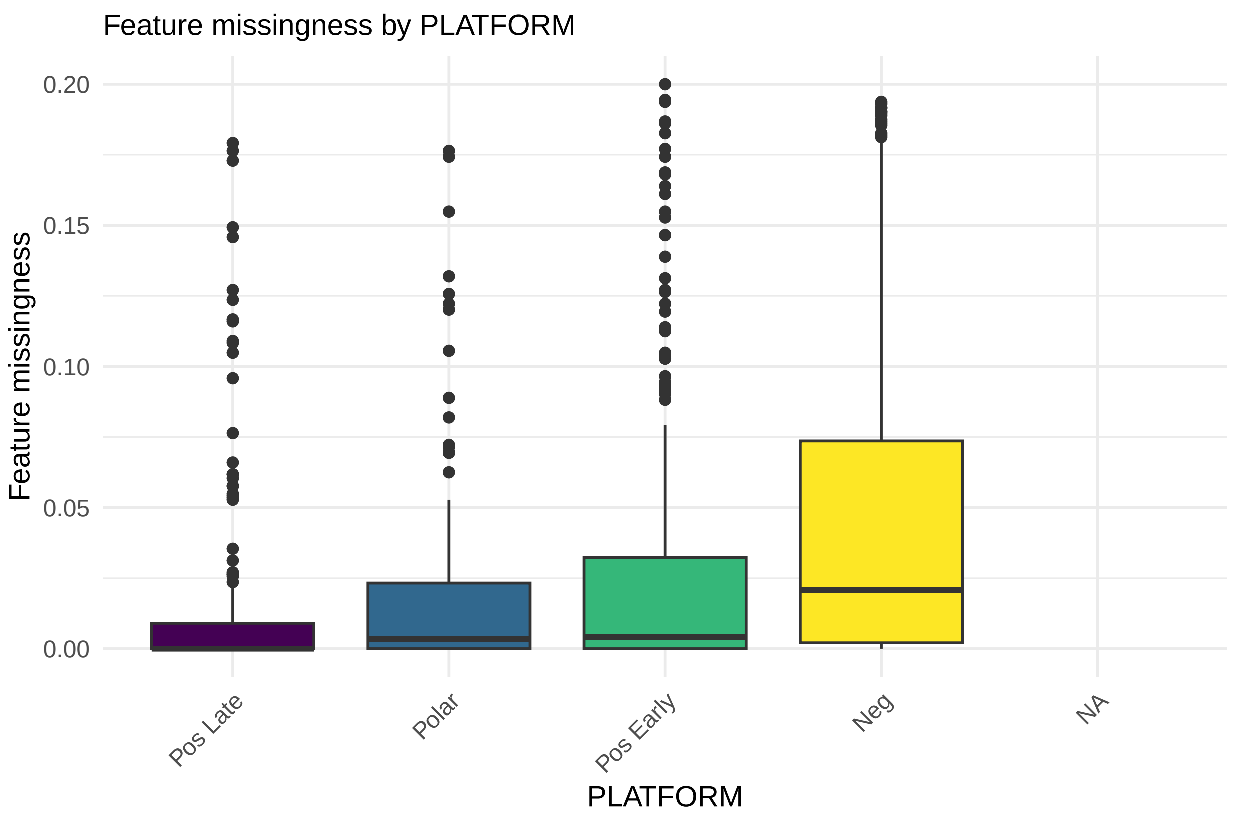

4.9% of the variation in Feature missingness can be explained by PLATFORM (pval = 2.27e−10)

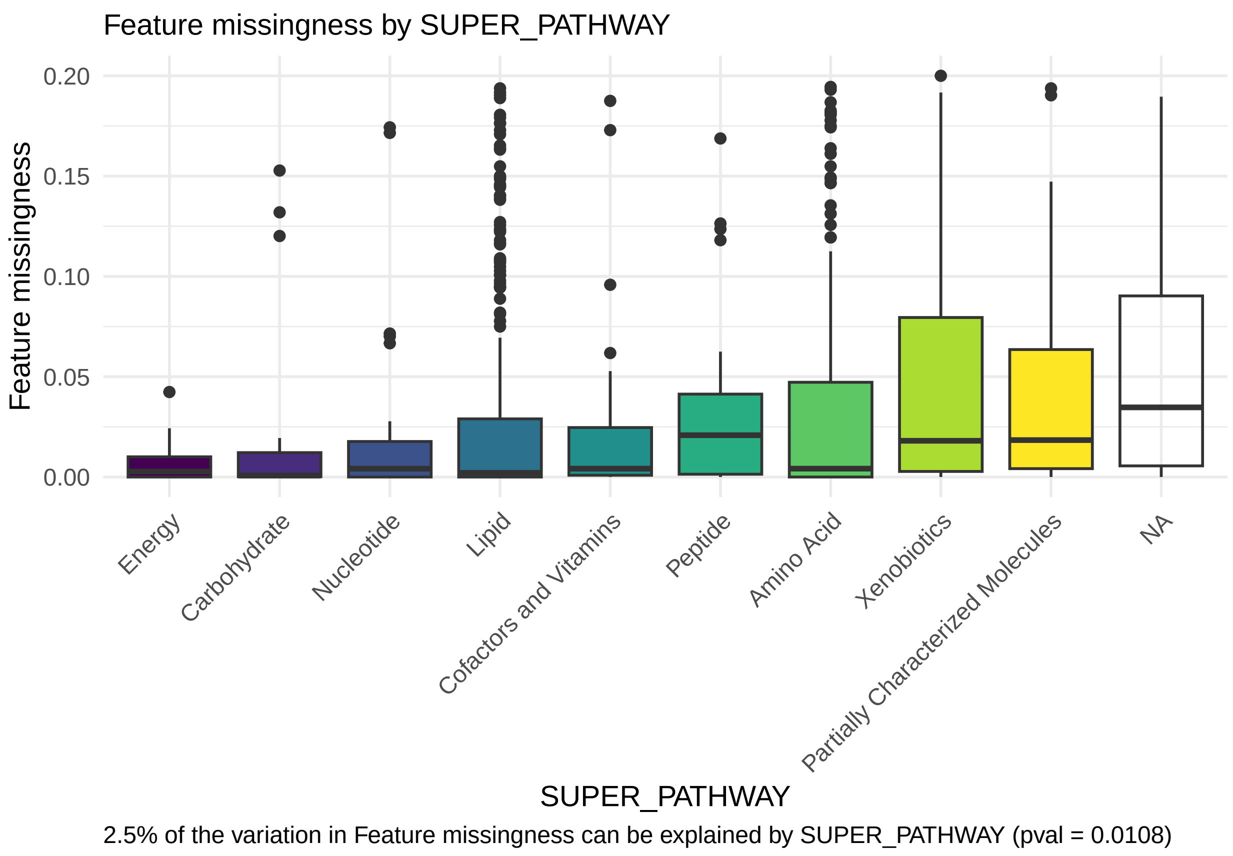
Feature missingness by SUPER_PATHWAY
Feature missingness
0.20
0.15
0.10
0.05
0.00
Energy
Carbohydrate
Nucleotide
Lipid
Cofactors and Vitamins
Peptide
Amino Acid
Xenobiotics
Partially Characterized Molecules
NA
SUPER_PATHWAY
2.5% of the variation in Feature missingness can be explained by SUPER_PATHWAY (pval = 0.0108)

Feature missingness by TYPE

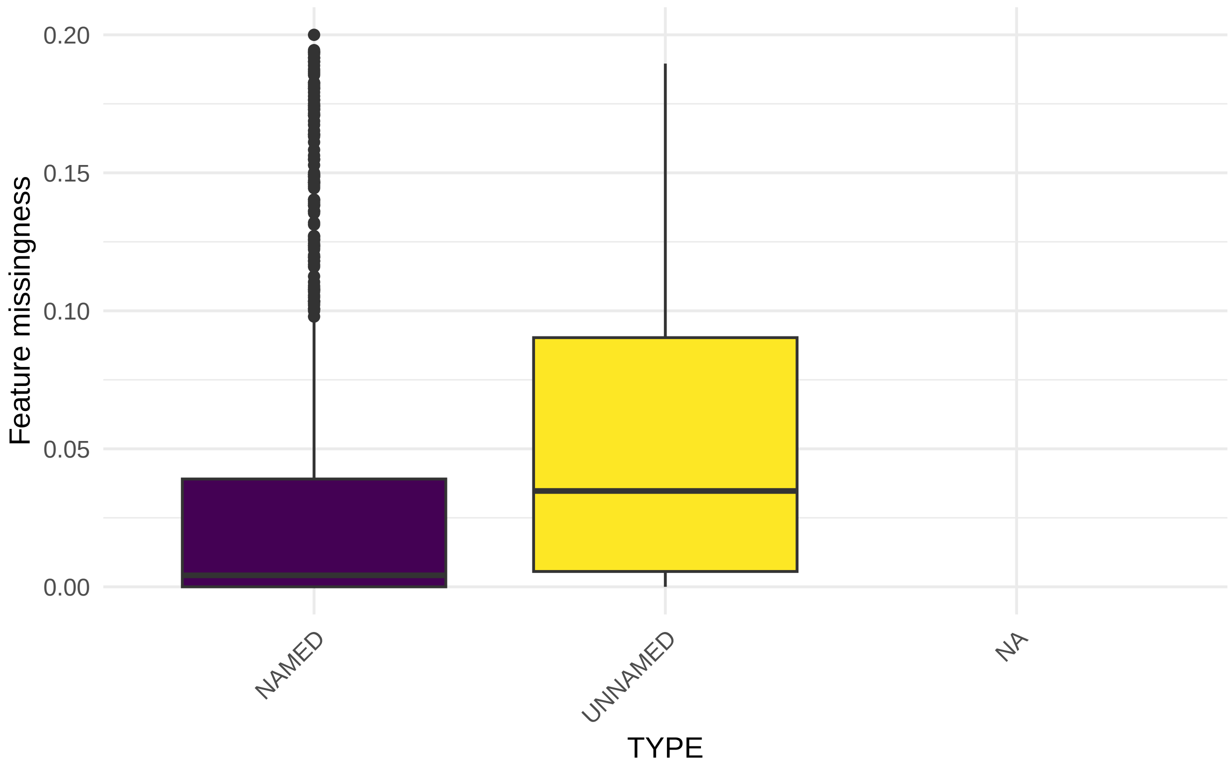

2.6% of the variation in Feature missingness can be explained by TYPE (pval = 4.64e−07)

#### 6.2 Sample missingness

The figure provides an illustrative evaluation of the proportion of *sample missingness* as a product of sample batch variables provided by your supplier. This is the univariate influence of batch effects on *sample missingness*. Box plot illustration(s) of the relationship that available batch variables have with sample missingness.

```
-- After filtering a total of 7 feature level batch variables were identified. --
-- They are:
   BOX_NUMBER
   GROUP_NAME
   NEG
   POLAR
   POS EARLY
   POS LATE
   WITHIN_SET_SAMPLE_NUMBER

-- After testing for redundancies a total of 3 feature level batch variables remain. --
-- They are:
   BOX_NUMBER
   GROUP_NAME
   WITHIN_SET_SAMPLE_NUMBER
```

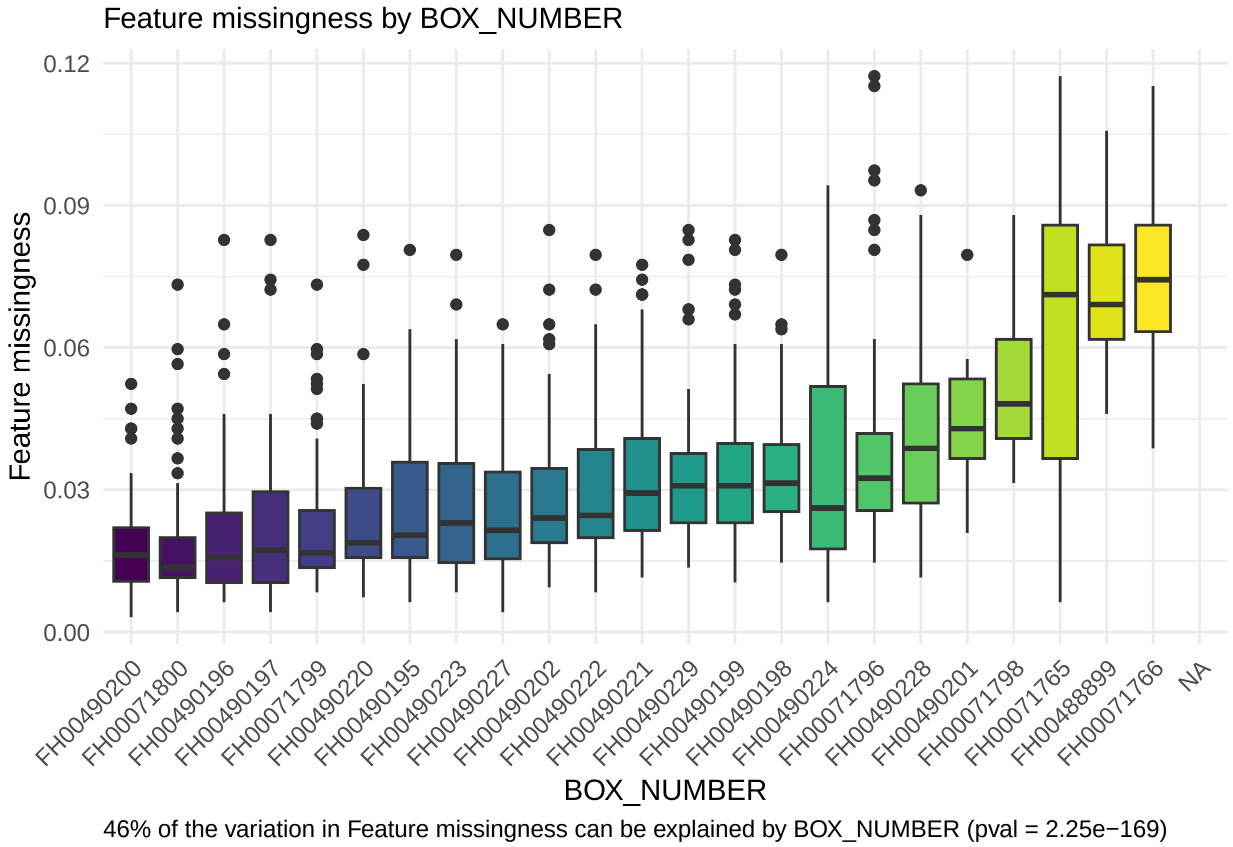

Feature missingness by GROUP_NAME

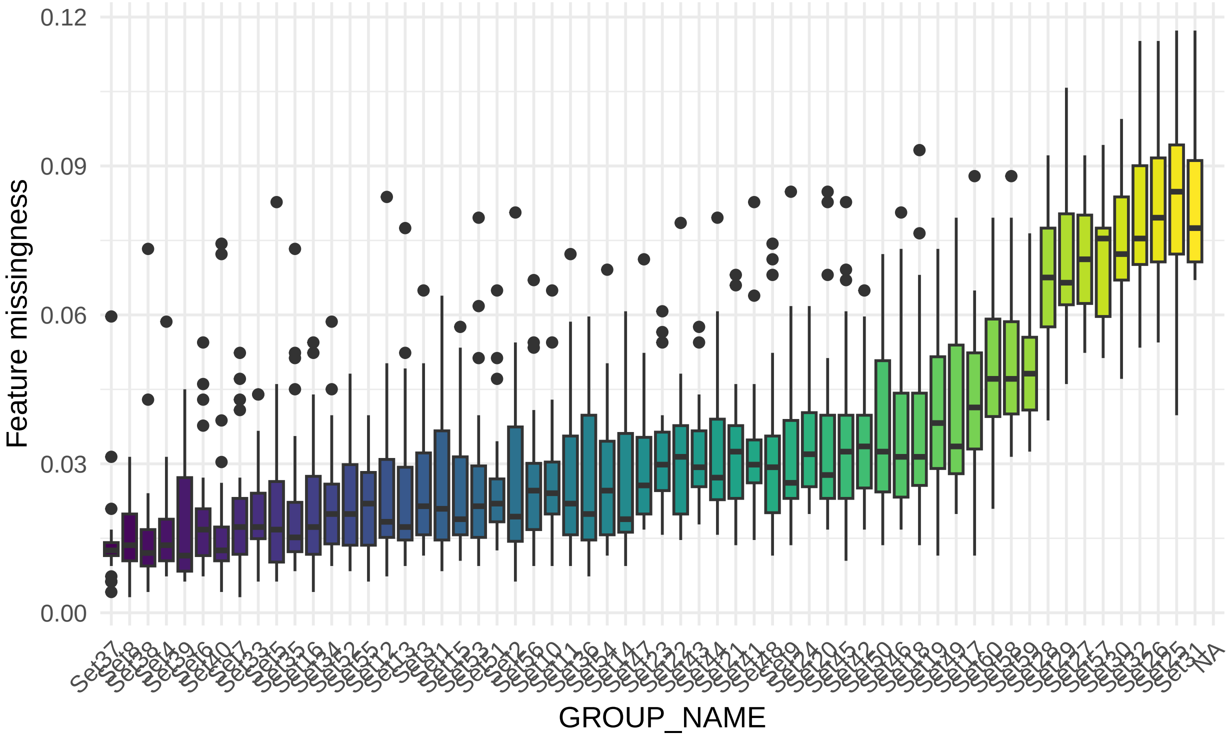

59% of the variation in Feature missingness can be explained by GROUP_NAME (pval = 9.99999999999999e−221)

Feature missingness by WITHIN_SET_SAMPLE_NUMBER

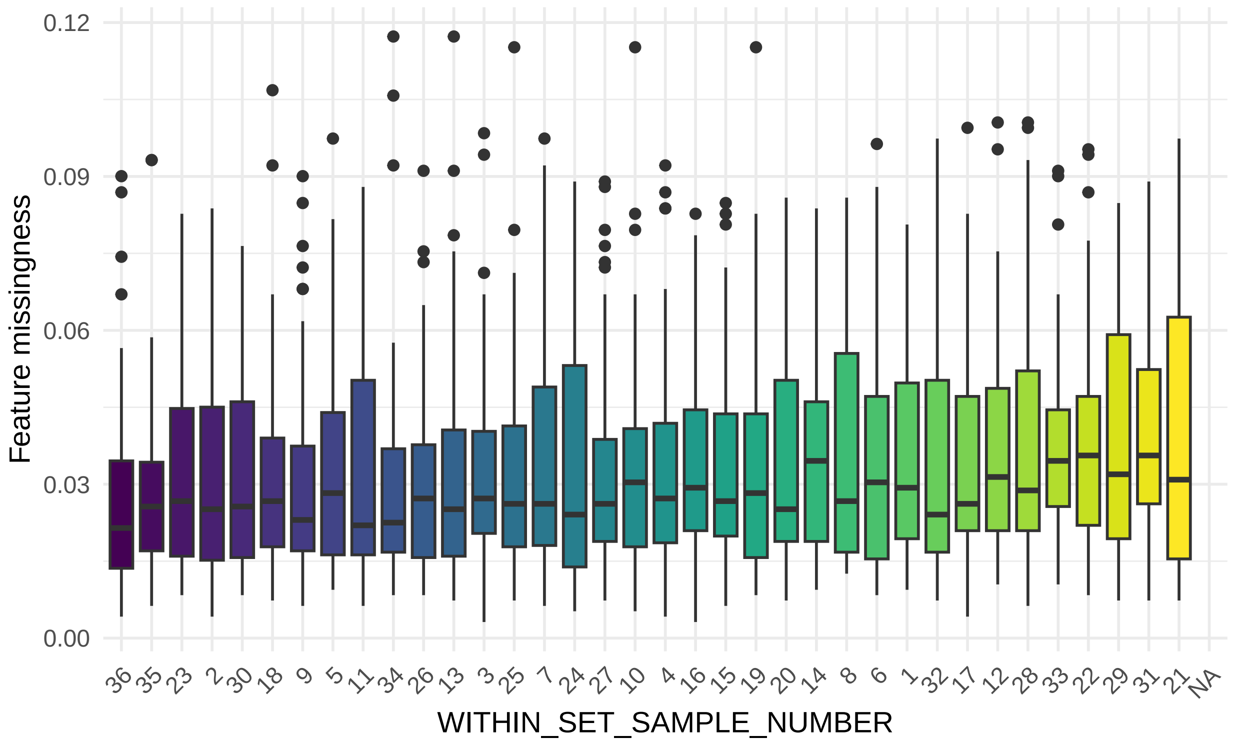

1.3% of the variation in Feature missingness can be explained by WITHIN_SET_SAMPLE_NUMBER (pval [illegible]
0.993)

#### 6.3 Multivariate evaluation: batch variables

TypeII ANOVA: the eta-squared (eta-sq) estimates are an estimation of the percent of variation explained by each independent variable, after accounting for all other variables, as derived from the sum of squares. This is a multivariate evaluation of batch variables on *sample missingness*. Presence of NA's would indicate that the model is inappropriate.

| batch.variable | etasq.var.exp | pvalue |
|---|---|---|
| box.number | 1.36 | 1.9388e−01 |
| group.name | 25 | 5.4934e−54 |
| within.set.sample.number | 2.13 | 2.7999e−01 |
| residuals | 71.51 | NA |

### 7 Total peak or abundance area (TA) of samples:

The total peak or abundance area (TA) is simply the sum of the abundances measured across all features. TA is one measure that can be used to identify unusual samples given their entire profile. However, the level of missingness in a sample may influence TA. To account for this we:

1. Evaluate the correlation between TA estimates across all features with PA measured using only those features with complete data (no missingness).
2. Determine if the batch effects have a measurable impact on TA.

#### 7.1 Relationship with missingness

Correlation between total abundance (TA; at complete features) and missingness. Relationship between total peak area at complete features (x-axis) and sample missingness (y-axis).

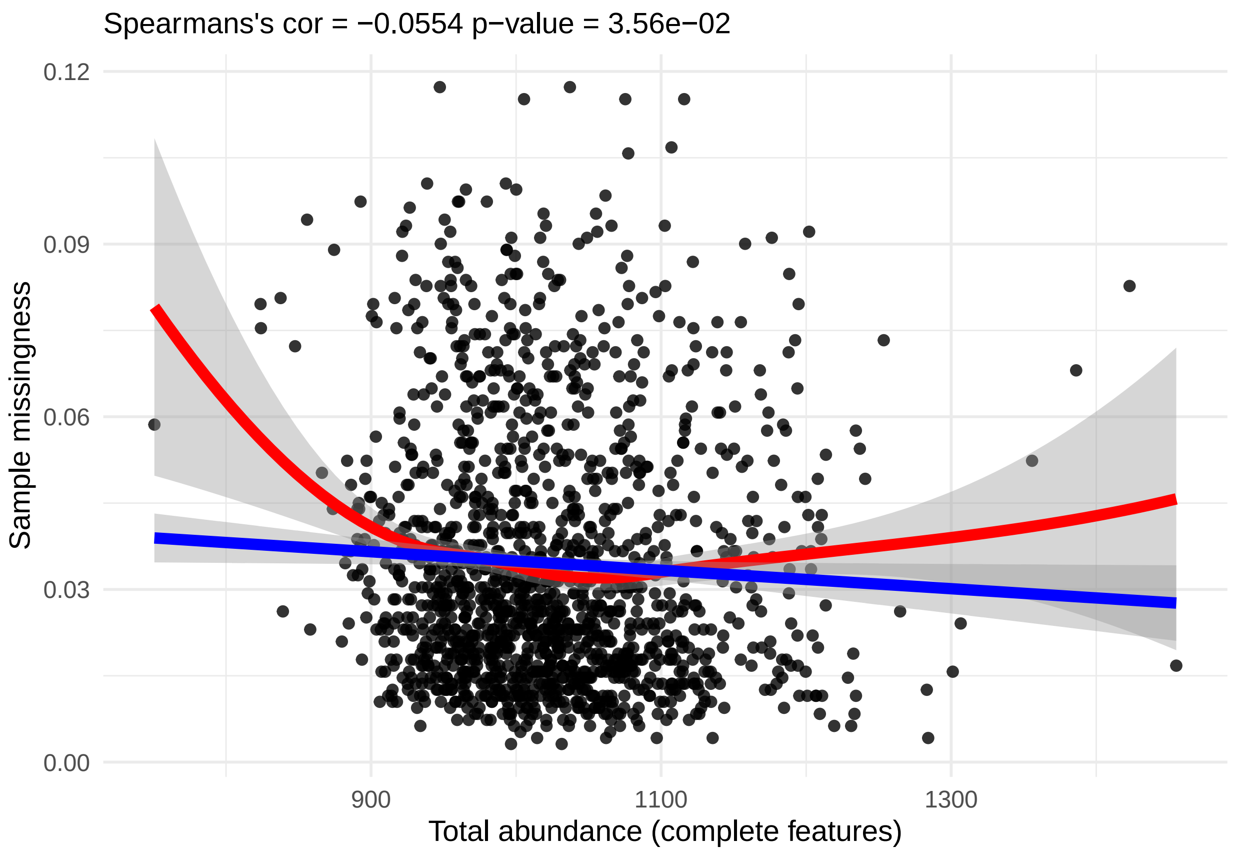

#### 7.2 Univariate evaluation: batch effects

The figure below provides an illustrative evaluation of the *total abundance* (at complete features) as a product of sample batch variables provided by your supplier. Violin plot illustration(s) of the relationship between total abundance (TA; at complete features) and sample batch variables that are available in your data.

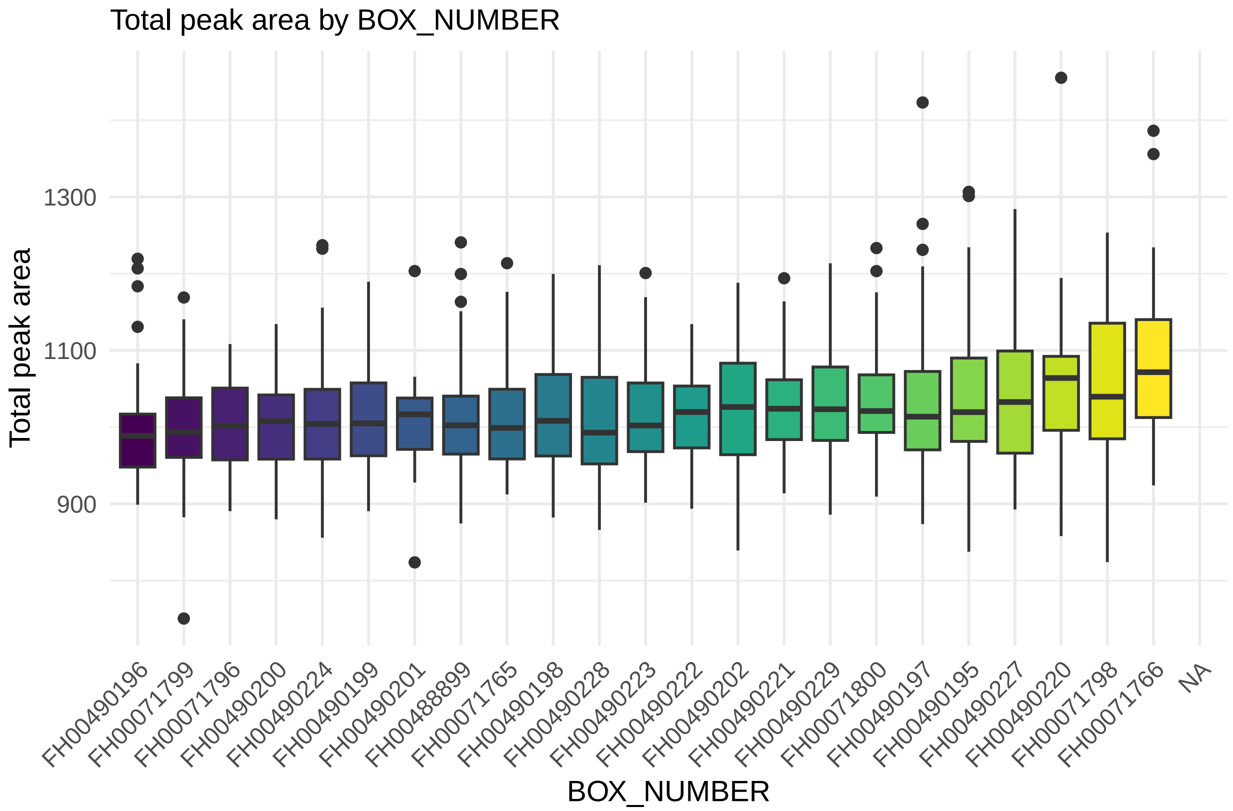

7.3% of the variation in Total peak area can be explained by BOX_NUMBER (pval = 3.2e−13)

Total peak area by GROUP_NAME

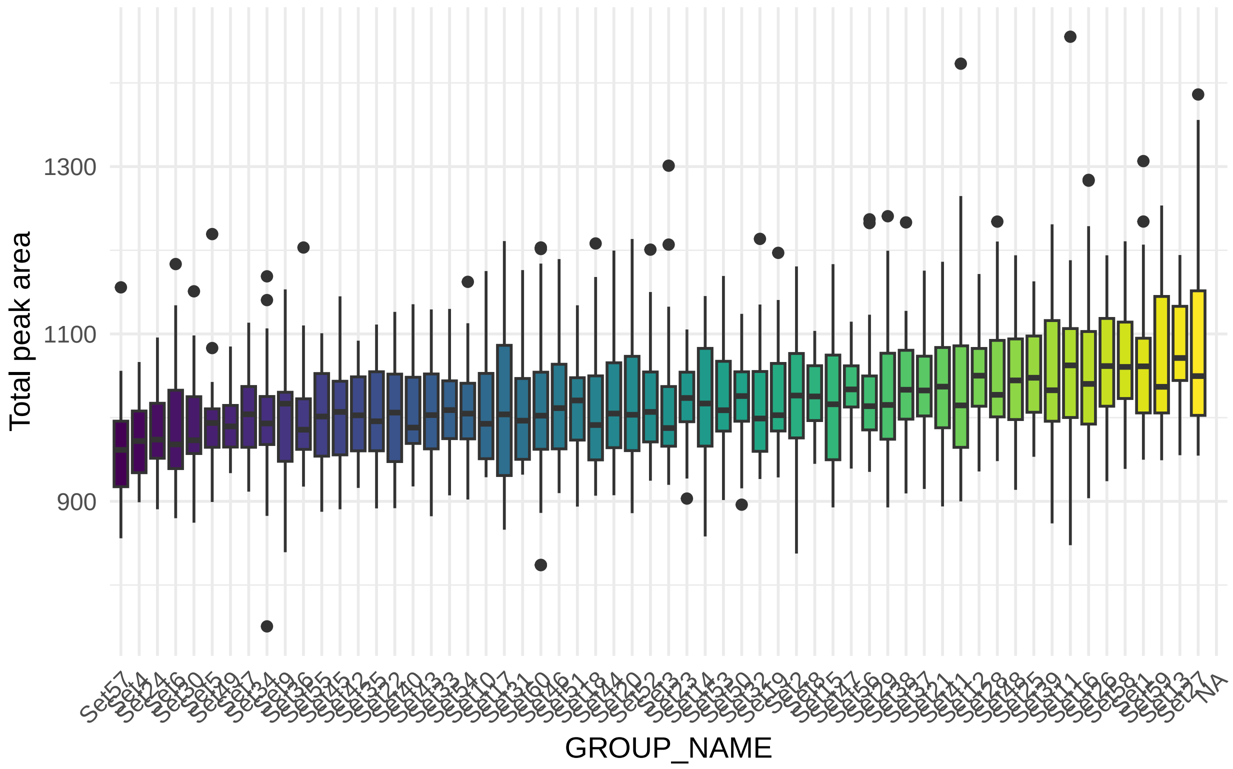

11% of the variation in Total peak area can be explained by GROUP_NAME (pval = 3.46e−12)

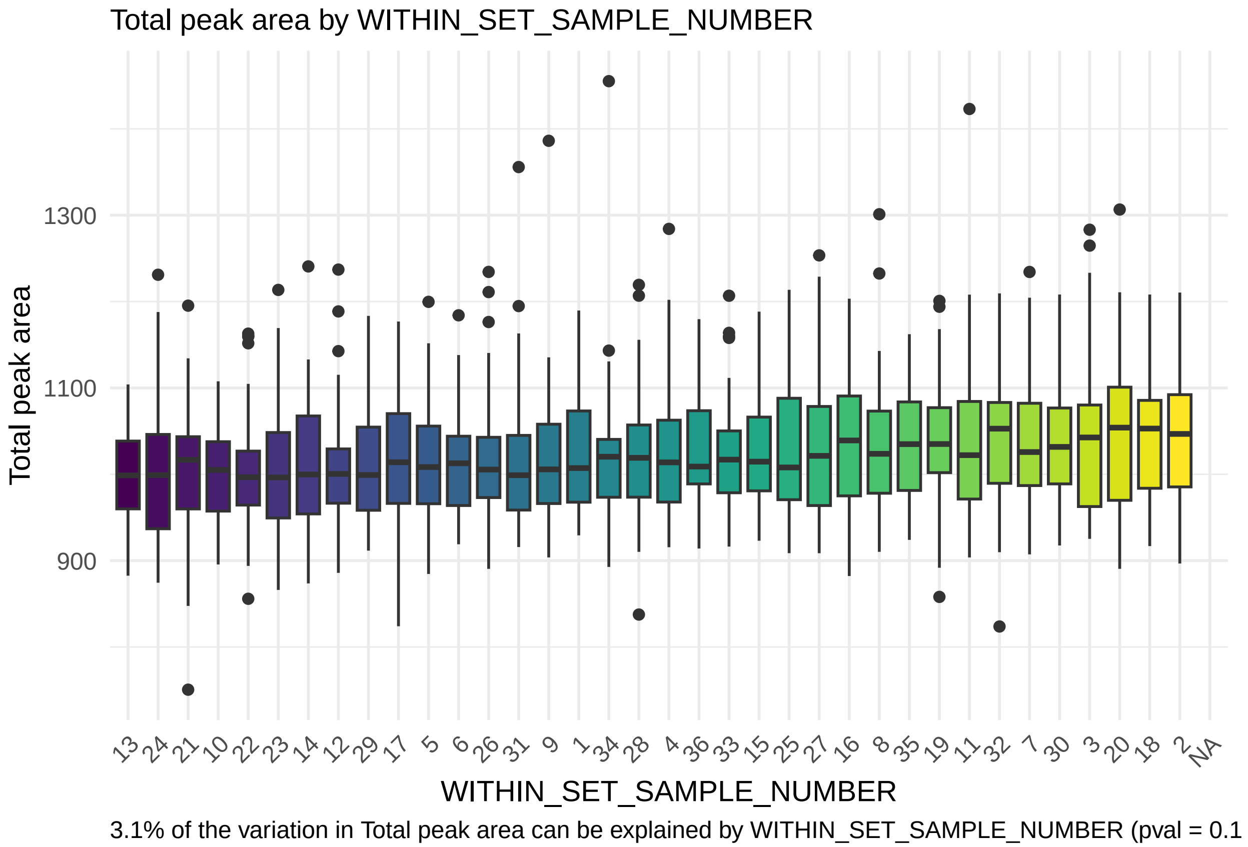

#### 7.3 Multivariate evaluation: batch variables

TypeII ANOVA: the eta-squared (eta-sq) estimates are an estimation on the percent of variation explained by each independent variable, after accounting for all other variables, as derived from the sum of squares. This is a multivariate evaluation of batch variables on *total peak|abundance area* at complete features.

| batch.variable | etasq.var.exp | pvalue |
|---|---|---|
| box.number | 1.72 | 1.8616e−01 |
| group.name | 5.82 | 8.5924e−03 |
| within.set.sample.number | 3.26 | 6.7865e−02 |
| residuals | 89.2 | NA |

#### 8 Power analysis

Exploration for case/control and continuous outcome data using the filtered data set

Analytical power analysis for both continuous and imbalanced presence/absence correlation analysis.

Simulated effect sizes (standardized by trait SD) are illustrated by their color in each figure. Figure (A) provides estimates of power for continuous traits with the total sample size on the x-axis and the estimated power on the y-axis. Figure (B) provides estimates of power for presence/absence (or binary) traits in an imbalanced design. The estimated power is on the y-axis. The total sample size is set to 99 and the x-axis depicts the number of individuals present (or absent) for the trait. The effects sizes illustrated here were chosen by running an initial set of simulations which identified effects sizes that would span a broad range of power estimates given the sample population's sample size.

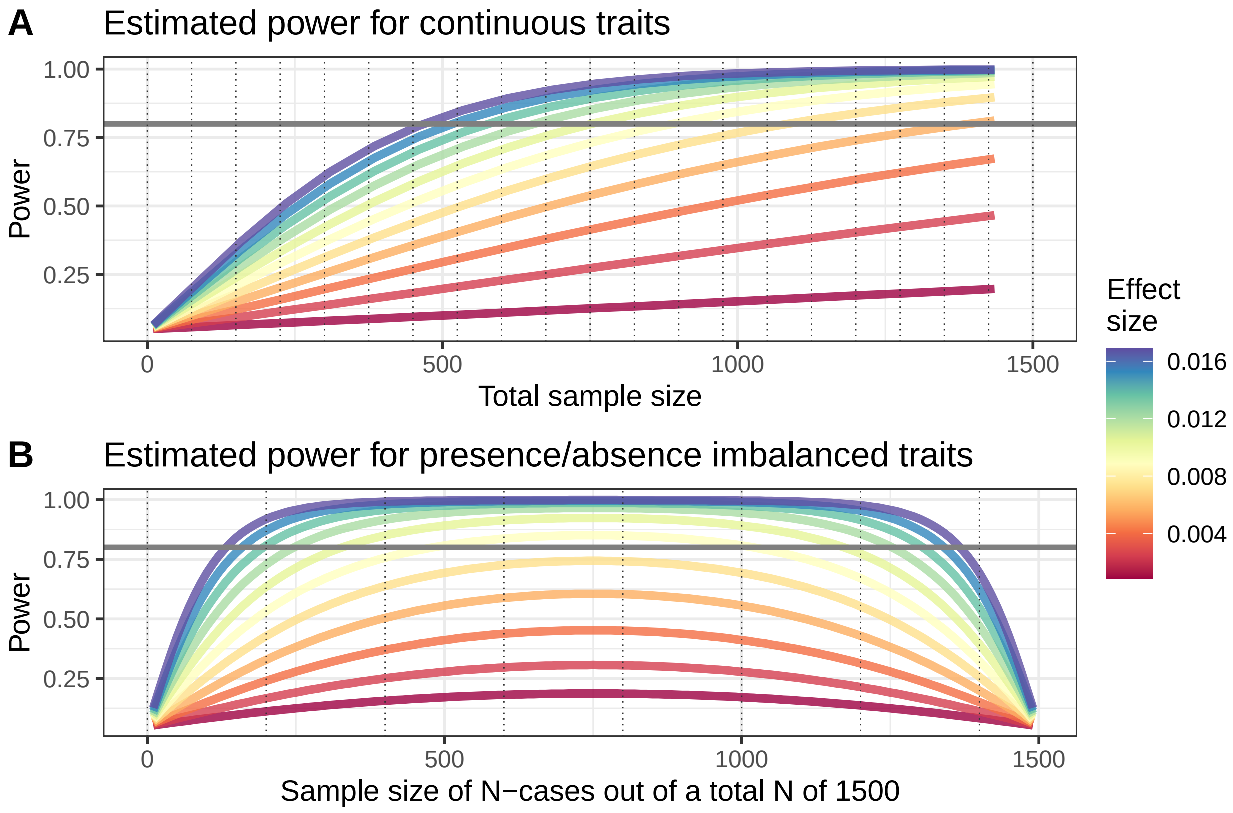
